# Resurvey of boreal earthworm communities in southern and northern Finland

**DOI:** 10.64898/2026.09.10.750052

**Authors:** Sylvain Gérard, Erin K. Cameron, Rebekka Kukowski, Alberto Piris, Helen R P Phillips

**Affiliations:** Department of Organismal and Evolutionary Biology, Faculty of Biological and Environmental Sciences, University of Helsinki, Helsinki, Finland; Department of Biology, Saint Mary’s University, Halifax, NS, Canada; Department of Biodiversity, Ecology and Evolution, Faculty of Biology, Universidad Complutense de Madrid, Spain

**Keywords:** Annelid, Conservation, Diversity, Subarctic, Assemblage

## Abstract

In northern boreal Europe, the Last Glaciation Maximum resulted in a tabula rasa of earthworm species, post-glacial recolonisation of few peregrine species. Boreal forests represent 11.5% of the Earth’s land area, and their cold winters represent the harshest season for earthworms. These biotopes are subject to substantial global changes, notably climate change and land use intensification, which might impact earthworm populations. Yet, no study measuring temporal changes in earthworm population or diversity is known in this biotope.

Here, we sampled earthworm communities in two northern and two southern areas in Finland forests. In total, 21 sites were surveyed in 2015 and resurveyed 10 years later in 2025-2026. We aimed at understanding how earthworm diversity and populations changed in the last 10 years, comparing community and species-specific diversity metrics.

In ten years, although not significant, earthworm abundance declined (−20%), whereas richness increased (+13%). The cold-tolerant *Dendrobaena octaedra* decreased, being the only one with a significant change. Temporal community and species dynamics followed a south-north gradient. Southern sites showed a decrease in abundance and *D. octaedra* populations, with an increase in abundance and occupancy of more heat-tolerant species. Northern sites showed an increase in abundance uniquely due to an increase in *D. octaedra* populations, but no other species were found there. These changes can be linked to climate changes leading to less rigorous winters in both south and north of Finland, and more rigorous summers in the south, potentially interacting with intense forest management, but could also be attributed to a still ongoing post-glacial recolonisation. These results call for more long-term sampling of earthworm communities in these fragile and rapidly changing environments.

## INTRODUCTION

The Last Glaciation Maximum, that ended 11,700 years ago, resulted in a *tabula rasa* of most earthworm populations in Northern Europe (Mathieu & Davies, 2014). Today’s European high latitude earthworm species pools are thought to be mainly the result of recolonisation from populations in southern regions that were not under ice sheets or permafrost, representing ecological refuges (Mathieu & Davies, 2014; Terhivuo, 1988). Some populations might also have survived in northern refuges, such as coastal areas or mountain summits emerging from ice sheets, and recolonise from there, as known as the nunatak hypothesis (Hansen et al. 2006; de Sosa et al. 2025; Stöp-Bowitz 1969). Such recolonisation is likely to have happened actively (earthworm active displacement) or passively (*via* streams, animal transport, or human activities) (Cameron et al., 2007; Terhivuo & Saura, 2006). It was therefore only possible for a restricted number of species showing high ability to disperse or to establish after passive dispersion. This somewhat recent biogeographical history resulted in northern European species pools being primarily composed of a small number of species, with good colonising abilities, usually named “peregrine species” by earthworm researchers (Gérard et al., 2025). Lumbricid species, native to southern and central Europe, are highly prevalent within this group of cosmopolitan species (Julin, 1949; Martinsson et al., 2021; Stöp-Bowitz, 1969; Terhivuo, 1988). This pattern is particularly striking in the Fennoscandian peninsula, where, as an example, Finland hosts just 16 species (Terhivuo, 1988) compared to *ca.* 200 species found in France (Gérard et al., 2025).

The Fennoscandian peninsula is characterised mostly by a boreal climate (*i.e.*, subarctic climate) (Beck et al., 2023), historically marked by short, cool summers and long, cold to very cold winters. These winters may represent a unique and intense environmental challenge for earthworms. In winter, earthworms face freezing temperatures that result in a lack of available water, typically frozen in surface soil layers, or at risk of having their body water frozen. Some individuals show frost avoidance strategies such as dispersion below the frost line, or frost tolerance, with accumulation of polyols in non-hatched individuals (*i.e.* individuals in cocoons) (Holmstrup & Zachariassen, 1996), or glucose in adults (Rasmussen & Holmstrup, 2002), resulting in the decrease of melting temperature of their body fluids (Holmstrup & Zachariassen, 1996). Indeed, differences in frost tolerance seem like an important driver in the distribution of species in Fennoscandia. Notably, in Finland *Dendrobaena octaedra,* a frost tolerant peregrine species, is widely distributed and particularly dominant in the north, whereas *Allolobophora chlorotica*, another peregrine species, showing lower frost tolerance, is only found in the southern coastal areas (Holmstrup & Zachariassen, 1996; Terhivuo, 1988).

Boreal forests have experienced increasing global change pressures in the last decades, such as intensification of forestry (in Fennoscandia, up to 90% of forest is managed (Gauthier et al., 2015)), fire regime shifts, or insect outbreaks (Gauthier et al., 2015). Climate change is also an important global change driver in boreal regions, with more intense droughts and a decrease in the number of days with subzero temperatures. Summers might therefore become a new, more widespread challenge for Fennoscandian earthworms. Indeed, in the rest of Europe, available water is lacking in summers and earthworms have to cope with water loss with seasonal dispersal (to wetter habitats or more in depth in soils), with cocoons laid before adults die, or through various forms of dormancy, *i.e.* ‘aestivation’ (quiescence, diapause) (Bouché, 1984). In this context of increasing pressures and changes in the course of seasons, associated with a still ongoing recolonisation of northern latitudes, we can expect that earthworm populations might change accordingly. Indeed, earthworm diversity is negatively impacted by global changes at global scale (Phillips et al., 2024), but there is low evidence on how it is impacted over time. In another boreal context, Cameron and Bayne (2015) found a rapid spread of invasive species in a resampling scheme of 6 to 7 years in Alberta, Canada. Spatially broader time-series analyses have found a decrease in earthworm abundance in the UK (Barnes et al., 2023, although this is debated; see Keith et al., 2026), or homogenization of communities in France (Gérard, 2024), although associated with an increase in local species richness. However, to date, no time-series analysis for earthworms in northern Europe are available.

As continuous sampling over time can be challenging, resampling sites previously studied can be a valuable method to understand how populations and diversity have changed due to global changes (Stuble et al., 2021). We sampled 21 sites in boreal forests in Finland, following a south-north gradient, in 2015 and again in 2025 or 2026. We use this resurvey to assess how earthworm community metrics (total abundance, richness, community composition) and earthworm species metrics (species abundance, occupancy) have changed over the last 10 years. We hypothesise that winter climate warming in Finland has accelerated the earthworm recolonisation of northern European boreal biomes. We would therefore expect a shift over time from earthworm communities dominated by cold-tolerant species to more diverse assemblages hosting more generalist species, particularly in the northernmost sites. However, this hypothesised richness increase is not necessarily expected to be accompanied with an increase in abundance, as other drivers of global change, such as forest management intensification, might restrict population sizes.

## MATERIALS & METHODS

### Study area and sampling

Sites studied in this work (n=21) are located in two areas of southern Finland (Tvärminne and Lammi) and in two areas of northern Finland (Levi and Kilpisjärvi) (Figure 1). These areas are designated as ‘regions’ in this work. In each region, 4 to 6 sites, spaced a mean of 14 km (sd=14km, min=0.26km, max=51km) apart, were sampled in 2015 and resampled in 2025 or 2026. The resampling took place at the same location (within *ca.* 5 meters) and at a similar time in the year (within a few days) as the historical sampling, minimizing intra-year seasonal variability in earthworm responses.

**Figure 1:**
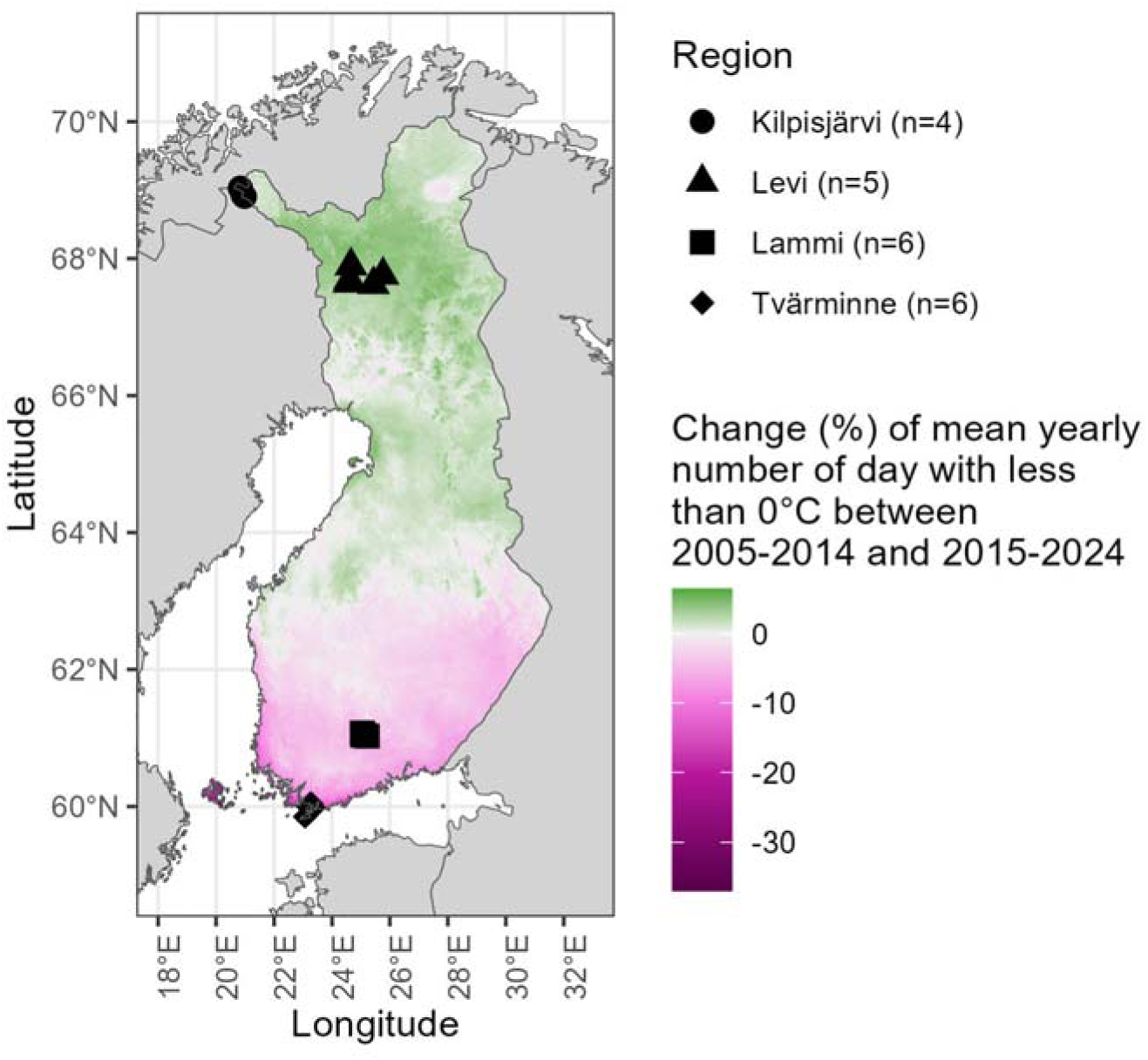
Location of studied sites in Finland. Country boundaries outlined in black. Sites studied in this work are represented as circles for Kilpisjärvi region, triangles for Levi region, squares for Lammi region and diamonds for Tvärminne region. Colours designate the % of change in mean number of days per year where daily temperature is less than 0 °C between 2005-2014 and 2015-2024 time periods, based on CHELSA data, for Finland. Other countries are shown in grey.

Tvärminne sites (n=6) are hemi-boreal forests, while Lammi (n=6), Levi (n=5), and Kilpisjärvi (n=4) sites are boreal forests. Lammi and Levi are found in a Taiga ecotone while Kilpisjärvi is found in a Taiga-Tundra ecotone. The habitats during the 2015 sampling consisted of forests dominated by spruce (*Picea abies*) (except Kilpisjärvi), birch (*Betula* spp.), or mixed forest (dominated by both *P. abies* and *Betula* spp.). One site in Kilpisjärvi area consisted of a young fir plantation (*Abies* sp.). Three sites experienced a clear cut between the initial sampling and the resampling (one in each of the most southern regions – ‘Tvärminne 8’, ‘Lammi 10’, ‘Levi 9’).

The same sampling protocol was applied in the historical sampling (2015) and the resurvey (2025-2026) (Figure 2). In each site, nine 25*25 cm quadrats were sampled along a 25 m transect. Six of the quadrats, which were spaced 5 m apart along the transect with alternate quadrats being placed 5 m perpendicular to the transect, were sampled by hand sorting the litter followed by mustard extraction. Two litres of mustard solution (10g of mustard powder per one litre of water) were applied, followed by 30 minutes of earthworm collection. Application of mustard solution consisted of a repetition of one-litre-application followed by 15 minutes of earthworm collection. The remaining three quadrats were sampled by hand sorting litter and soil to a depth of 20 cm. These quadrats were placed at 0, 10 and 20 m along the transect, 1 m perpendicular to the transect. All specimens found in mustard extraction and hand sorting were collected. Specimens were then killed and stored in ethanol (70% ethanol in 2015, 100% ethanol in 2025-2026). Ethanol was renewed within 7 days.

**Figure 2.**
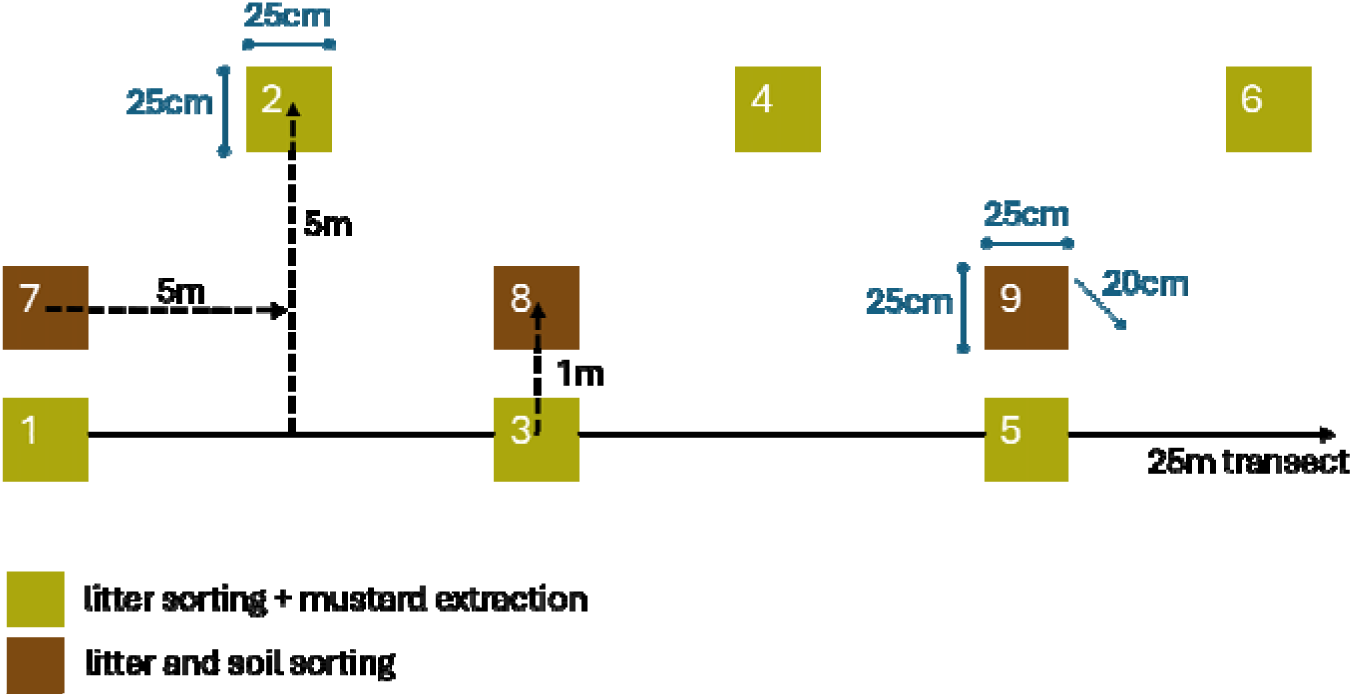
Sampling scheme conducted at each site. Quadrats numbers (1 to 9) are the same as in the data used for this work.

### Species identification

Species identification was conducted on ethanol-preserved specimens, within 3 months of sampling. Each individual was identified with a microscope to the best taxonomic resolution possible. Specimens sampled in 2015 were identified by EC using Reynolds et al. (1977) and Sherlock (2012). Specimens sampled in 2025-2026 were identified by SG using Terhivuo (1988) (for the list of species found in Finland) and Sherlock (2012) (for the morphological traits used for identification). We harmonised scientific names based on Brown et al. (2023). Specimens were considered as ‘adult’ when they showed a clitellum, others were considered as juvenile. Juveniles without developed reproductive traits were identified using the key in Table 1. This key is solely based on Terhivuo (1988) Finnish species list, as no adult specimen of species not corresponding to this list were found in both sampling events.

**Table 1:** Key for identifying juvenile earthworm specimens in the sites sampled for this work.

|  |  |  |
| --- | --- | --- |
| 0. | no pigmentation | 1. |
|  | pigmentation | 2. |
| 1. | setae widely paired posteriorly | ***Octolasion*** sp.<sup>a</sup> |
|  | setae closely paired throughout all body | Undetermined<sup>b</sup> |
| 2. | setae closely paired | 3. |
|  | setae distant (or widely paired) | 5. |
| 3. | tanylobous prostomium | ***Lumbricus*** sp.<sup>c</sup> |
|  | epilobous prostomium | 4. |
| 4. | dark brown pigmentation, body length can exceed 8 mm | ***Aporrectodea longa***<sup>d</sup> |
|  | reddish pigmentation (can be quite light), body length always less than 8 mm. | ***Eiseniella tetraedra***<sup>e</sup> |
| 5. | epilobous prostomium (sometimes almost tanylobous), purplish color in preserved specimen, antero-posterior pigmentation gradient less pronounced | ***Dendrobaena octaedra***<sup>f</sup> |
|  | epilobous prostomium, reddish color in preserved specimen, antero-posterior pigmentation gradient more pronounced | ***Bimastos rubidus***<sup>f</sup> |

<sup>a</sup> Possible species: *O. cyaneum*, *O. lacteum*. <sup>b</sup> Possible species: *Aporrectodea* *caliginosa*, *Ap. rosea*, *Allolobophora chlorotica*. <sup>c</sup> Possible species: *L. terrestris*, *L.* *rubellus*, *L. castaneus*, *L. festivus*. <sup>d</sup> Possible confusion with *Ap. trapezoides*, but it has not been recorded in Finland. <sup>e</sup> Also matches *Eisenia* spp. (*E. fetida*, *E. andrei*) and *Dendrobaena hortensis*, but in Finland these species are always associated with "leaf composts, manure heaps, greenhouses, waste, soils etc., i.e. sites with the constant addition of organic material to the soil by human" (adapted from Terhivuo, 1988). <sup>f</sup> Both could be mistaken with *Satchellius mammalis* found in neighbouring
countries, but there is no known record in Finland, and the male pore is usually visible in juveniles.

### Climate change data

We extracted from CHELSA-daily dataset (Karger, 2025) daily mean near-surface air temperature (‘tas’) for each day between 01 January 2005 to 31 December 2024 and thus for the whole area of Finland (Figure 1, Figure S1. To calculate the number of freezing days, the number of days for each year with a daily mean near-surface air temperature of less than 0°C was calculated and averaged for the 2005-2014 and 2015-2024 time periods. The rationale for choosing these time periods was to have representative climate data for each sampling event, so we selected a 10-year span preceding each sampling event. Climate data from 2020 were removed because of NA values in some days of the year.

We also extracted climate data from the four Finnish Meteorological Institute (FMI) meteorological stations that were the closest to sampled sites (Table S1, Figure S2). We extracted daily average, minimum, and maximum temperatures (‘tday’, ‘tmin’ and ‘tmax’) as well as snow depth (‘snow’) from 01 January 2005 to 31 December 2024 from FMI open data project (Honkola et al. 2013). We used the same method to measure the number of freezing days, using the ‘tday’ variable. We also averaged these variables for the 2005-2014 and 2015-2024 time periods.

Data extraction was conducted using R packages Rchelsa (https://gitlabext.wsl.ch/karger/rchelsa) and fmi2 (Lehtomäki & Lahti, 2026). Geographic data were manipulated using R packages *terra* (Hijmans et al., 2026) and *sf* (Pebesma et al., 2026) and were mapped using R packages *tidyverse* (Wickham & RStudio, 2023), *scales* (Wickham et al. 2025), *tidyterra* (Hernangómez et al., 2026), and *rnaturalearth* (Massicotte et al., 2026). All computation and figures were done using R version 4.5.2 (2025-10-31 ucrt) (R Core Team, 2020).

### Analyses

We computed community and species-specific metrics for both 2015 and 2025-2026 sampling events. For species-specific metrics, we measured abundance and occupancy (*i.e.* sum of occurrence) for each species. For community metrics, we measured community abundance and species richness. Undetermined pieces of earthworm with no anterior part were not considered in the measurement of abundance, and only individuals identified to species were retained in the calculation of richness.

As our sampling effort is somewhat small (n=21 sites), and metrics did not follow a normal distribution, we used Wilcoxon signed rank tests to compare community metrics between 2015 and 2025-2026. To compare species abundance between time periods, we performed a one-tailed paired t-test for abundance of each species observed in 2015 and in 2025-2026, with 999 permutations. This test was developed for resurvey context by Legendre (2019).

We also computed temporal beta diversity metrics for each site, accounting for abundance. We used Legendre (2019) index “TBI” (Temporal Beta diversity Index) that corresponds to Bray-Curtis dissimilarity, and associated indices “TBI gains” and “TBI losses”, corresponding to components of TBI associated to, respectively, gains and losses of species occurrence and abundance.

Data manipulation and figures were done using R and *tidyverse* R package. One-tailed paired t-tests and TBI indices computation were done using R package *adespatial* (Dray et al., 2026).

Data and code used in this work are available in GitHub (**link, to be provided**) and Zenodo (**doi, to be provided**). Data will also be available in GBIF.

## RESULTS

### Change in climate

CHELSA data show that in the last 10 years, southern Finland (latitude <63°N), where the Tvärminne and Lammi sites are located, has experienced a decrease in number of freezing days compared to the previous 10 years (Figure 1). In contrast, northern Finland (latitude >63°N), including the Kilpisjärvi and Levi sites, has experienced an increase in the number of freezing days overall (Figure 1). The observed increase showed a smaller amplitude than the observed decrease, with a maximum increase of +6.5% of freezing days and a maximal decrease of -36.9% of freezing days. Southwestern Finnish coasts showed the highest decrease in number of freezing days. The whole country exhibited an increase in mean temperature between the two time periods (Figure S1).

Data from the meteorological stations of the FMI that are the closest to our sites show a decrease in snow depth in southern sites and an increase in northern sites (Table S1). All mean, minimum and maximum temperatures increased, with the largest increase observed in southern sites (+0.66°C in Tvärminne and +0.31°C in Kilpisjärvi in 10 years). Number of freezing days showed the same south-north gradient than CHELSA data, with a decrease in southern sites (−24.0 days in Tvärminne, -8.1 days in Lammi), but an increase only for the northernmost region (−0.3 days in Levi, +1.0 day in Kilpisjärvi).

### Change in species specific metrics

In total, eight species were found (Figure 3A), all belonging to Lumbricidae. *Lumbricus castaneus* was only found during the resampling in one Tvärminne site (Figure 3B), and all seven other species were found in both sampling events (Table S2). *Dendrobaena octaedra* was the only species found in Kilpisjärvi sites for both sampling events, and in Levi sites in the resampling (Figure 3B, Table S2). Species abundance showed high variability (high standard deviation, typically larger than mean) (Table S2).

**Figure 3:**
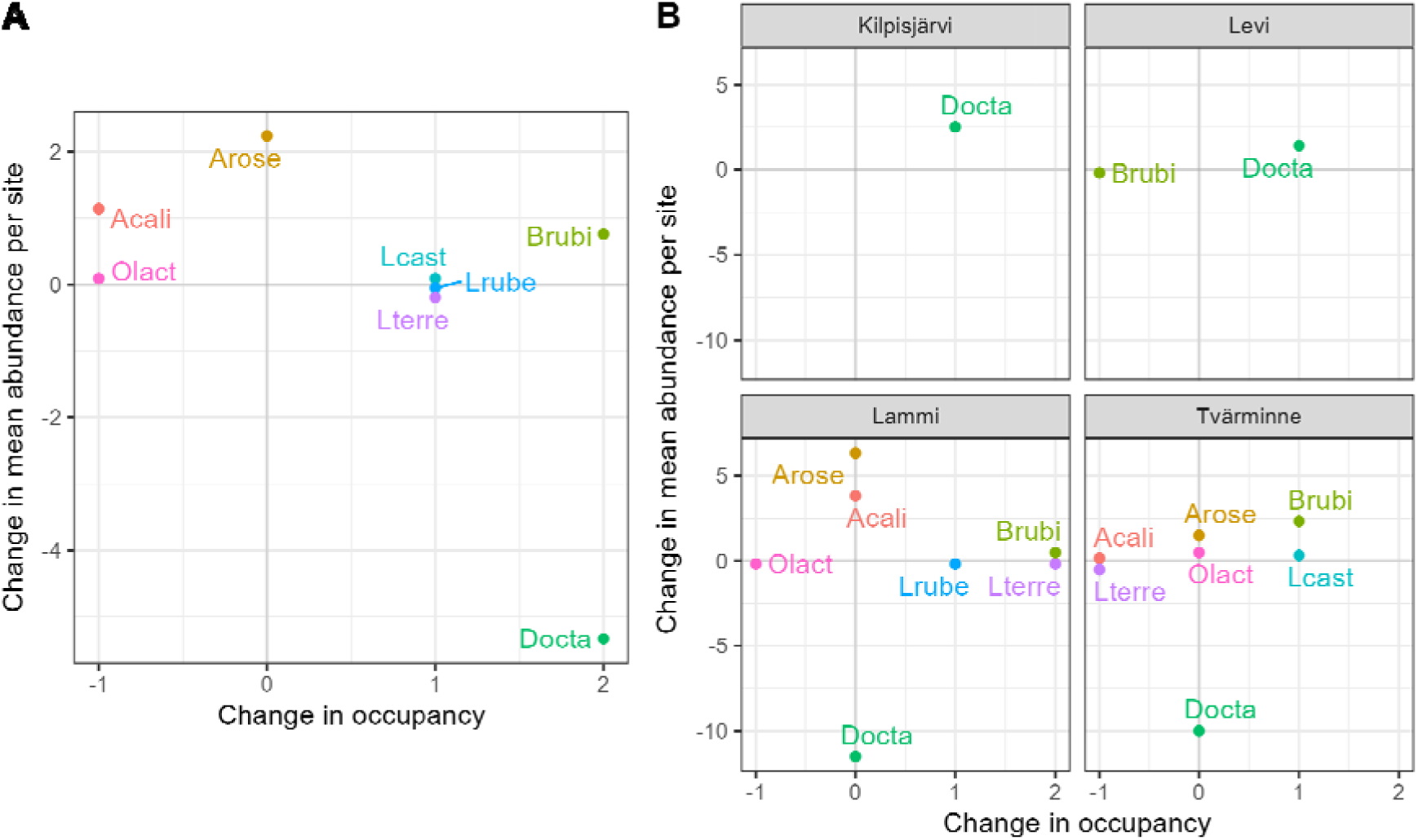
Change in abundance per site and occupancy of species found in the survey and the resurvey for all sites (A) and among each region (B). In each site 0.5625 m² were sampled (0.375 m² for litter sorting + mustard extraction, 0.1875 m² for litter and soil hand sorting). List of species plotted and their acronyms in parenthesis: *Aporrectodea caliginosa* (Acali), *Aporrectodea rosea* (Arose), *Bimastos rubidus* (Brubi), *Dendrobaena octaedra* (Docta), *Lumbricus castaneus* (Lcast), *Lumbricus rubellus* (Lrube), *Lumbricus terrestris* (Lterre), *Octolasion lacteum* (Olact).

*Dendrobaena octaedra* was the only species showing a significant change in abundance per site, with a significant decrease. The other species showed no significant change in abundance. *Aporrectodea rosea* had the highest positive value of abundance change (Figure 3A). The abundance decrease in *L. terrestris* was mostly driven by a single site in Lammi, where individuals were reduced from 12 to 5.

In terms of occupancy, *Bimastos rubidus* had the highest occupancy increase. *D. octaedra* showed the highest occupancy change, matching the numbers shown by *B. rubidus*, but lost occupancy when considering adults only (Table S2). Increase in occupancy was more frequent than decrease (5 *vs.*2 species)*. A. rosea* was the only species where occupancy stayed the same between the initial sampling and the resampling.

In southern sites (Lammi and Tvärminne regions), *D. octaedra* showed the larger decrease in abundance, but no change in occupancy (Figure 3B). Other species had smaller abundance changes, and some species had consistent increases in abundance such as *A. caliginosa*, *A. rosea* or *B. rubidus*. More species had a positive occupancy change in Lammi, compared to Tvärminne where occupancy increase and decrease where balanced. In northern sites (Levi and Kilpisjärvi regions), *D. octaedra* showed both increase in abundance and in occupancy, and *B. rubidus* was only found in one site in Levi in the initial sampling, without being found in the resampling (Figure 3B).

### Change in population metrics

#### Abundance

There was no significant change in abundance per site between the initial sampling and the resampling, but abundance of adults in the Tvärminne sites showed a near-significant decrease (P-value = 0.05791) (Table 2, Table S3). For all individuals, abundance numbers were smaller in the resampling overall (−20%), which was mainly driven by sites in the southernmost region (Tvärminne sites, -48%). The two northernmost regions studied, Kilpisjärvi and Levi sites, had higher abundances in the resampling (respectively +333% and +75%).

**Table 2:**
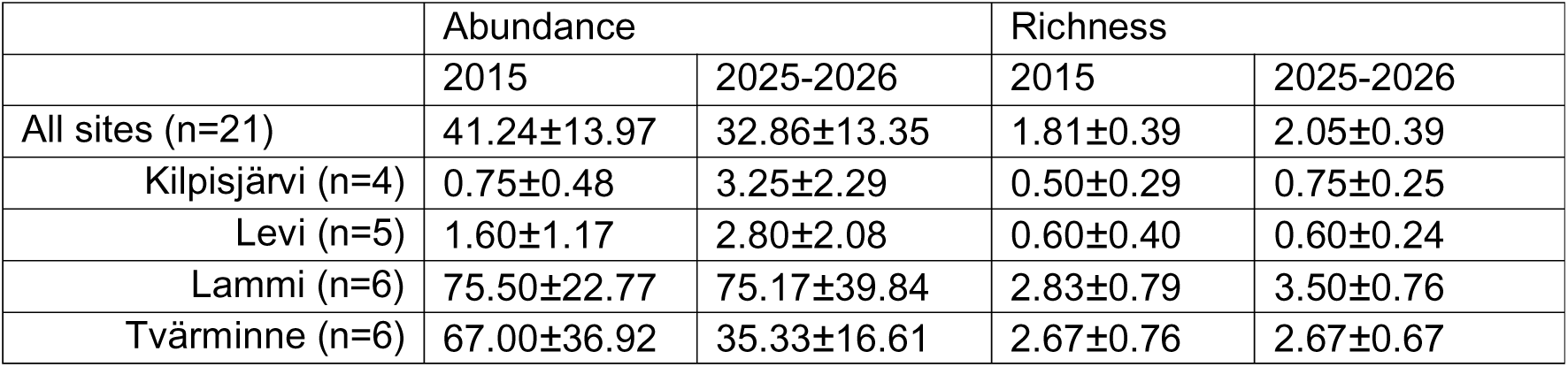
Earthworm abundance and species richness per site in the initial sampling (2015) and the resampling (2025-2026). In each site 0.5625 m² were sampled (0.375 m² for litter sorting + mustard extraction, 0.1875 m² for litter and soil hand sorting). Results are showed as mean ± standard deviation. The table shows mean site abundance and richness for all sites and among each region. P-value of Wilcoxon signed rank test all > 0.1.

|  | Abundance |  | Richness |  |
| --- | --- | --- | --- | --- |
|  | 2015 | 2025-2026 | 2015 | 2025-2026 |
| All sites (n=21) | 41.24±13.97 | 32.86±13.35 | 1.81±0.39 | 2.05±0.39 |
| Kilpisjärvi (n=4) | 0.75±0.48 | 3.25±2.29 | 0.50±0.29 | 0.75±0.25 |
| Levi (n=5) | 1.60±1.17 | 2.80±2.08 | 0.60±0.40 | 0.60±0.24 |
| Lammi (n=6) | 75.50±22.77 | 75.17±39.84 | 2.83±0.79 | 3.50±0.76 |
| Tvärminne (n=6) | 67.00±36.92 | 35.33±16.61 | 2.67±0.76 | 2.67±0.67 |

Three sites have experienced a clear cut between the initial sampling and the resampling, one site in the Levi, Lammi and Tvärminne regions. The site in the Levi region showed no individuals in both sampling events, however, the sites in Lammi and Tvärminne showed large decreases in the number of individuals found (clear cut site in Lammi: 17 individuals in 2015 *vs.* 2 individuals in 2025; clear cut site in Tvärminne: 17 individuals in 2015 *vs.* 4 individuals in 2026).

Two sites in Levi region had no individuals in both sampling events. Two sites in Kilpisjärvi and one in Levi regions had no individual in the initial sampling, but at least one in the resampling. One site in Kilpisjärvi region had one individual in the initial sampling but none in the resampling.

#### Species richness

There was no significant change in richness between the initial sampling and the resampling (Table 2, Table S3). Richness numbers were higher in the resampling (+13%), due to higher richness in Kilpisjärvi and Lammi in the resampling (+50% and +24%, respectively). As communities in Kilpisjärvi sites are composed of only *D. octaedra*, higher richness measured in the resampling there was due to local colonization of *D. octaedra* in sites previously empty.

#### Temporal change in communities

Temporal beta diversity (TBI) per site averaged 0.575±0.294 (Figure 4). Its component explained by gains (TBI gains) and losses (TBI losses) were close, averaging respectively 0.314±0.364 and 0.260±0.317. Kilpisjärvi sites exhibited the highest TBI (0.917), greatly explained by high TBI gains, *i.e.,* a gain in species abundance. Levi sites showed less TBI (0.471) but were almost entirely explained by gains in species abundance. Southern sites, in Lammi and Tvärminne regions, had TBI of same range as Levi sites (respectively 0.537 and 0.437), but had their larger TBI components in the loss of species (occupancy or abundance), particularly for Tvärminne sites. It should be noted that a temporal change from no individual to more than one individual, identified to species level, will result in the highest possible TBI of 1.

**Figure 4:**
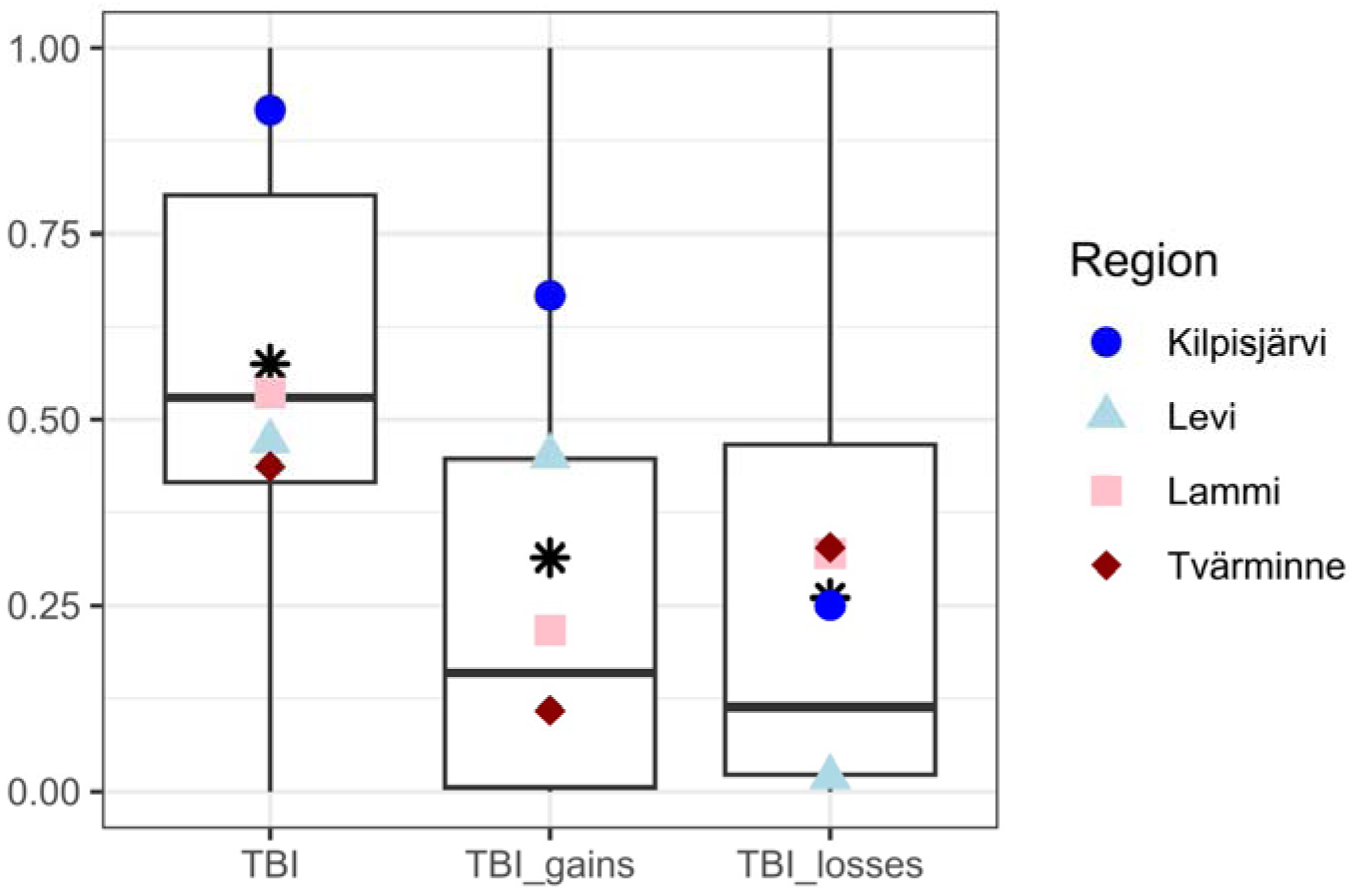
Change in species composition (or ‘temporal beta diversity’) between the initial sampling and the resampling, measured with various indicators. ‘TBI’ (*i.e.*, Temporal Beta-Diversity Index, corresponding to Bray-Curtis dissimilarity), ‘TBI_gains’ and ‘TBI_losses’ were calculated with function TBI of *adespatial* R package (Legendre, 2019) and measure changes in species composition taking into account abundances. ‘TBI_gains’ represents the part of TBI explained by gain of species locally, while ‘TBI_losses’ represents the part of TBI explained by loss of species locally. Boxplot representation with black asterisk, dark blue circle, light blue triangle, pink square, and dark red diamond as means of, respectively, all sites, Kilpisjärvi region, Levi region, Lammi region, Tvärminne region.

## DISCUSSION

### Overall change in community metrics

Our results show that earthworm abundance appears to be decreasing. This apparent abundance decrease is coherent with Phillips et al. (2024) metanalysis showing that global changes negatively impact earthworm diversity (including abundance). It is also consistent with Barnes et al. (2023) claim that earthworm populations are undergoing a massive decline in the UK, but Keith at al. (2026) contradicted these results and showed no change after reanalysing the same data. While statistical tests are likely not significant due to a lack of statistical power for such an effect size, we also stress the importance of not over-emphasizing the potential decline until additional data is collected. Indeed, time-series data on earthworm communities is rare, and collecting such data should be a priority.

Richness does not seem to decrease, in fact showing a small increase, but as again, it is not significant we cannot accept our initial hypothesis of a richness increase. However, in a 60-year earthworm resampling scheme distributed across France, Gérard (2024, Chapter 3A) did find a significant increase in local richness. Typically, no change in local richness as a response to global change is likely the result of local extinction of species being balanced by the local introduction of species more adapted to the new conditions (Dornelas et al., 2019, 2023; Vellend et al., 2013). Local colonization events can also predate the local extinction events, resulting in an early richness increase (Kuczynski et al., 2023). In our study, there is currently no clear indication of local extinction of species, but increasing colonisation within regions may be underway. Indeed, one of the main criticisms of time-series analyses, particularly those that show no richness change, is the length of the time-series, with shorter time-series being unable to adequately detect changes (Gonzalez et al., 2016).

### Species – specific responses

*Dendrobaena octaedra* was the only species showing a significant change in abundance, with a large overall decrease (despite having a small occupancy increase). This species is the most common species in Finland (Terhivuo, 1988), primarily due to being frost tolerant and having parthenogenetic reproduction, both adaptations suitable for colonizing the Finnish environment (Holmstrup & Zachariassen, 1996; Terhivuo & Saura, 2006). While this species shows an overall decrease, it shows opposing dynamics within the different regions in our study. Its populations appear larger in northern regions whereas the biggest declines are seen in the southernmost regions. This suggests that northern regions might be becoming more suitable for *D. octaedra*, even though it is already adapted to cold climates, in opposition to southern regions.

The overwhelming dominance of *D. octaedra* among Finnish earthworm fauna in boreal forests contributed to its own dynamics driving most of the community abundance dynamics observed. In northern regions (Kilpisjärvi and Levi), except one occurrence of *Bimastos rubidus* in Levi in the initial sampling, it is the only species found, and it is the most abundant in southern regions (Lammi and Tvärminne). As this is also one of the few species that could be identified to species level at all life stages, the results for this species are likely more robust. Additionally, as some of the other species can show cryptic diversity (e.g., *Aporrectodea rosea* or *Lumbricus terrestris*) which is not visible with our morphology-based identifications, this might make observed changes more intricate.

Other species were less widely distributed and therefore had lower occupancy, resulting in potentially no detectable change. However, among them, occupancy increase was more frequent than occupancy decrease. Locally, this could be again explained by local colonization preceding the longer process of local extinctions in changing environments (Kuczinski 2023). The two species with highest abundance increase (*A. caliginosa*, *A. rosea*) and the only species not found in the initial sampling but found in the resampling (*L. castaneus*) are species that are restricted to southern Finland and Åland archipelago, especially the latest two (Nieminen et al., 2011; Terhivuo, 1988). The decrease of *L. terrestris,* mostly due to one site in Lammi with 12 *vs.* 5 individuals found, is less easily interpretable.

### South-North gradient response

Community and species-specific metrics showed a south-north gradient in their temporal responses. Abundance increased in the north but decreased in the south. *D. octaedra* populations increased as well in the north and decreased in the south, while species that usually occur more in the south saw their populations increase in the south, with no colonisation of the north. However, richness change did not respond across this south-north gradient.

We initially hypothesised that climate change would lead to a shift from earthworm communities dominated by cold-tolerant species to more diverse communities, hosting generalist species, especially in the north. Our data partially support this hypothesis, but notably only in the south. Indeed, *D. octaedra*, the species most tolerant to frost, declined in Tvärminne and Lammi, whereas species more tolerant to heat – and therefore likely less tolerant to frost – such as *A. rosea*, *A. caliginosa* and *L. castaneus* are increasing. Thus, *D. octaedra* dominance is shifting to more heat-tolerant species dominance. However, in the north (Kilpisjärvi and Levi), the species more tolerant to heat did not appear, with one species (*B. rubidus*) not re-sampled, and the frost-tolerant *D. octaedra* increasing. It could be explained by an intermediate situation, where changes in environmental conditions are beneficial to *D. octaedra*, but these sites are still not suitable for species more adapted to southern conditions, or that these species have yet to colonise.

Meteorological stations (Honkola et al. 2013) and modelled climate (Karger, 2025) both show that the south of Finland is experiencing a change in climate conditions. Notably, frost and snow are decreasing at unprecedent rates, with temperatures increasing. Less harsh winters, but also drier and hotter summers might be beneficial for heat-tolerant species, thus summer drought might be becoming a stronger filter for species than winter. On the other hand, in the north, climate change might lead to less rigorous winters, reducing the intensity of winter harshness filter, without necessarily having summers acting like a strong filter, explaining why a species like *D. octaedra* is thriving there. However, climate variables like snow depth or frost did not show a decrease but seem stagnant in the north. Other climatic factors such as decrease in snow season length (Aalto et al. 2026) could be potential explanations of seemingly better conditions for *D. octaedra*, whereas increase in rain-on-snow events (Aalto et al. 2026), thus decreasing snow insulation capacity, could mitigate this.

Another notable anthropic pressure occurring across Finland is forest management. In Finland, forests are actively logged and old-growth forests decline is alarming (Kortesalmi et al. 2025). Intense forest management is now considered as the primary threat to biodiversity in Finland (Hyvärinen et al. 2019; Kortesalmi et al. 2025). Clear forest cuts can have negative impacts on soil fauna, particularly on macroarthropods and decomposers, and these negative impacts are still seen after 10 years (Siira-Pietikäinen & Haimi 2009). Three sites in our study had experienced a clear cut during the ten-year span between sampling, and two of those showed a large decline in earthworm abundance, with the third one having no individuals in either sampling event. Therefore, forest management could potentially act as a strong driver of earthworm decline and interact with the negative effect of the changing climate. Other land use changes, e.g., transition from boreal forests to mines (Gololobova & Legostaeva 2023; Lassila 2021), urbanisation, or outbreaks of forest pests (Lyytikäinen-Saarenmaa & Tomppo 2002; Neuvonen & Viiri 2017) might also have important negative impacts in Finland, at least locally.

### Anthropogenic-driven changes or post-glaciation recolonisation?

We associated observed changes to anthropogenic drivers such as climate change and land use intensification to biodiversity change. However, the north of Europe has been experiencing a post-glaciation recolonisation of earthworms in the last 11,000 years. Most of this recolonisation is thought to be due to human-induced introductions from southern European refuges. In Finland, this is likely to have happened in the last few centuries, as the most northern parts of Finland were only permanently settled in the 18th century (Terhivuo, 1988). These post-glacial recolonisations might also originate from northern ice-free refuges, that likely existed in Fennoscandia during the last glaciation (Hansen et al. 2006, de Sosa et al. 2025). We can therefore legitimately wonder whether the observed change should be attributed to recent anthropogenic changes or a post-glacial recolonisation still ongoing.

Parts of North America were largely covered by ice during the last glaciation as well. These areas also experienced post-glacial recolonisations, yet, mostly originating from European settlers introducing lumbricids from European populations as early as the 16th century (Gates 1982). Thus, in the *tabula rasa* hypothesis context, boreal forests from Finland and North America have somewhat experienced a similar process, i.e. a colonisation of previously glaciated areas by similar lumbricid species likely introduced by human activities in a similar time frame (the last few centuries). In the *nunatak* hypothesis, this earthworm recolonisation should be much older, but could have been facilitated in the last century by human activities. In both cases, as Fennoscandia and north America experienced a somewhat analogous post-glacial process, and recolonisations are still ongoing in north America (some parts have been colonised only recently, Mathieu et al. (2024); Scheu & Parkinson (1994)), the change we observe might possibly be a post-glacial recolonisation stage, not necessarily a recent, anthropogenic-induced change of communities that had previously reached a steady state. Indeed, our results showing a larger number of occupancy increases compared to decreases, as well as *D. octaedra* experiencing an increase of both abundance and occupancy in the north of Finland, despite these areas not exhibiting the larger climate changes, are evidence for species in the process of colonisation. Climate change could act as a catalyst of colonisation of *D. octaedra* in the north and other, heat-tolerant, species in the south, but might hinder success of already well-established *D. octadrea* populations in the south.

### Limits of the study design

We acknowledge that our study is based on a relatively small number of sites, implying small statistical power. Therefore, our observations and conclusions rely on trends. However, we argue that our results are coherent with the literature. The time scale of this study is also rather small, which might underestimate long term anthropogenic-driven changes, and the biggest changes might have happened in the preceding decades (Mihoub et al. 2017). However, re-sampling for earthworms is rare, and a 10-year gap is rather large in earthworm research and a common threshold for biodiversity change assessments (van Klink et al. 2020). Also, this work is based on resampling, a method that is sometimes criticised, as it is highly dependent on inter-year variability, potentially leading to erroneous conclusions (Stuble et al., 2021). Earthworms therefore experience high inter-, but also intra-year variability (Schmidt & Curry 2001). Inter year variability can either over-or under-estimate temporal changes (Boennec et al. 2024) in the hypothesis of a linear change. Intra-year variability is less likely to impact our result as we matched the sampling dates between both sampling events. However, as time series are severely lacking for earthworms, we argue that it is important to take opportunities that exist, like resurveys of past data, even recent ones, to try to capture how these essential organisms are impacted by global changes. Therefore, we call here for more long-term sampling of earthworm communities throughout boreal forest but also globally.

## CONCLUSION

Earthworms are essential organisms in areas where they are not invasive, promoting crops and acting as ecosystem engineers (Blouin et al. 2013; Fonte et al. 2023). There is now growing evidence that they are impacted by changing environments, pleading for better protection but also for more temporal assessments in diverse conditions (biomes, land uses, areas hosting variable proportions of rare and endemic species). This will be necessary to fully understand how human impact them and what we can expect for them in the future. There is an urging need to start applying spatially broad, long-term earthworm sampling schemes, on top of taking opportunities of resampling old data, that can reveal substantial insights into temporal dynamics of earthworm populations.

## Acknowledgements

We want to acknowledge Sámi people and their ancestral lands, in which we sampled for this work. We also want to thank the University of Helsinki for a free of charge stay in Tvärminne research station. This work was possible thanks to the Research Council of Finland (Grant number 362759).

