## Supplementary Material for "Resurvey of boreal earthworm communities in southern and northern Finland"

**APPENDIX**

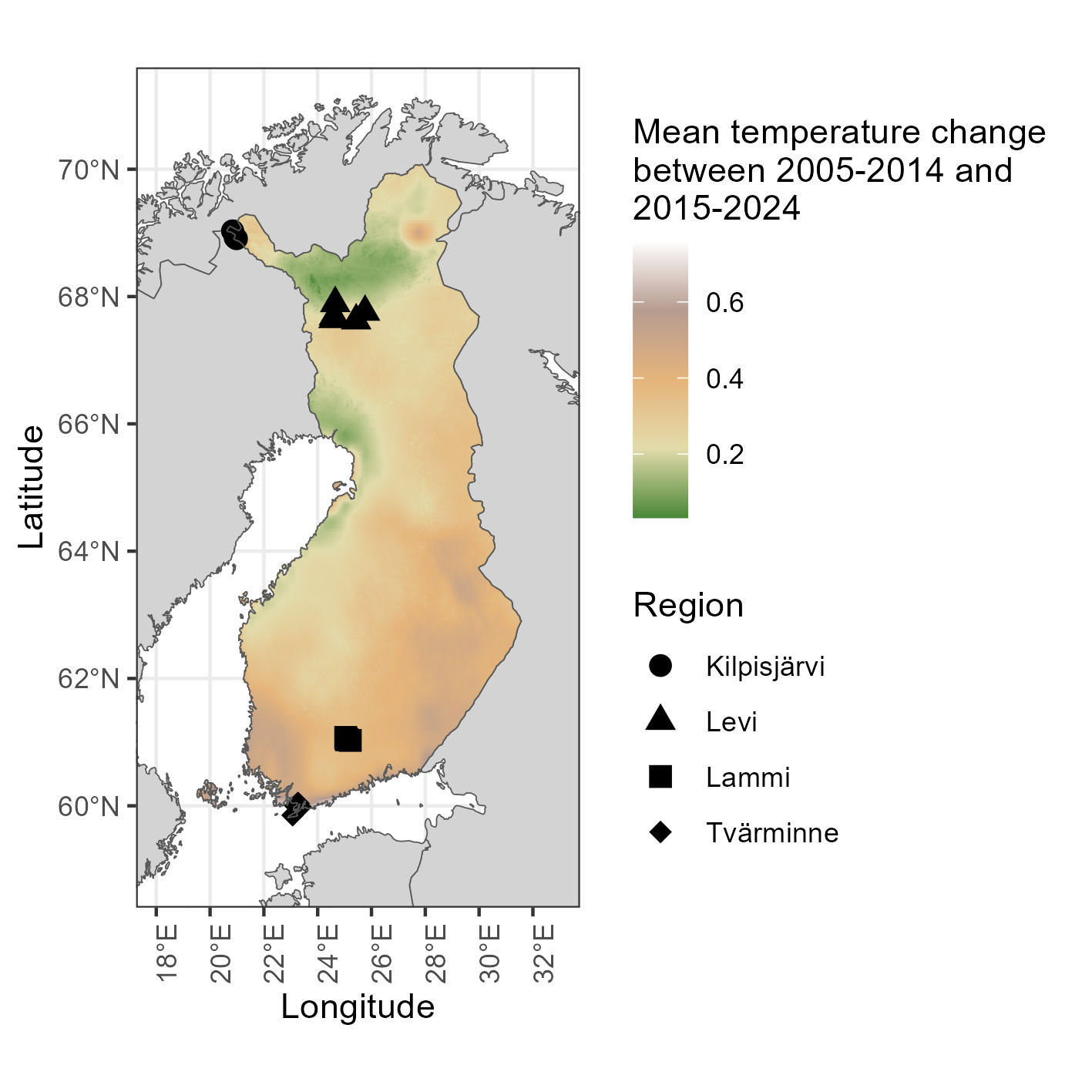

Figure S1: Change in mean temperature (°C) in Finland between 2005-2014 and 2015-2024 time periods, based on CHELSA data. Country boundaries outlined in black. Sites studied in this work are represented as circles for Kilpisjärvi region, triangles for Levi region, squares for Lammi region and diamonds for Tvärminne region. Other countries are shown in grey.

Table S1: Climate variables extracted from meteorological stations of the Finnish Meteorological Institute (FMI) from 01 January 2005 to 31 December 2014 (‘2005-2014’ in the table) and from 01 January 2015 to 31 December 2024 (‘2015-2024’ in the table). Kilpisjärvi data extracted from station ‘Enontekiö Kilpisjärvi’ (FMISID: 102016, Latitude: 69.0, Longitude: 20.8); Levi data extracted from station ‘Sodankylä Tähtelä’ (FMISID: 101932, Latitude: 67.4, Longitude: 26.6); Lammi data extracted from station ‘Hämeenlinna Lammi Pappila’ (FMISID: 101154, Latitude: 61.1, Longitude: 25.0); Tvärminne data extracted from station ‘Hanko Tvärminne’ (FMISID: 100953, Latitude: 59.8, Longitude: 23.2). ‘Snow depth’, ‘Mean temperature’, ‘Min temperature’ and ‘Max temperature’ correspond to average daily values for the period considered of variables ‘snow’, ‘tday’, ‘tmin’ and ‘tmax’ of available variables of meteorological stations of FMI. ‘Number of freezing days’ corresponds to the average number of days in the year with a mean temperature (‘tday’) of less than 0°C for the period considered.

|  | **Snow depth (cm)** | | **Mean temperature (°C)** | | **Min temperature (°C)** | | **Max temperature (°C)** | | **Number of freezing days** | |
| --- | --- | --- | --- | --- | --- | --- | --- | --- | --- | --- |
|  | **2005-2014** | **2015-2024** | **2005-2014** | **2015-2024** | **2005-2014** | **2015-2024** | **2005-2014** | **2015-2024** | **2005-2014** | **2015-2024** |
| Kilpisjärvi | 31.409328 | 35.246626 | -1.0284193 | -0.7174776 | -5.0322925 | -4.836295 | 2.989611 | 3.297840 | 185.2 | 187.0 |
| Levi | 25.543701 | 29.916218 | 0.7420477 | 1.1681243 | -3.7991204 | -3.391605 | 5.125364 | 5.413704 | 165.1 | 164.6 |
| Lammi | 8.582627 | 7.421376 | 4.9156637 | 5.5205196 | 0.9399788 | 1.563092 | 8.938167 | 9.457326 | 109.1 | 100.3 |
| Tvärminne | 4.994721 | 1.704993 | 6.6535407 | 7.3177766 | 3.9083132 | 4.636403 | 9.434412 | 9.979365 | 75.5 | 57.4 |

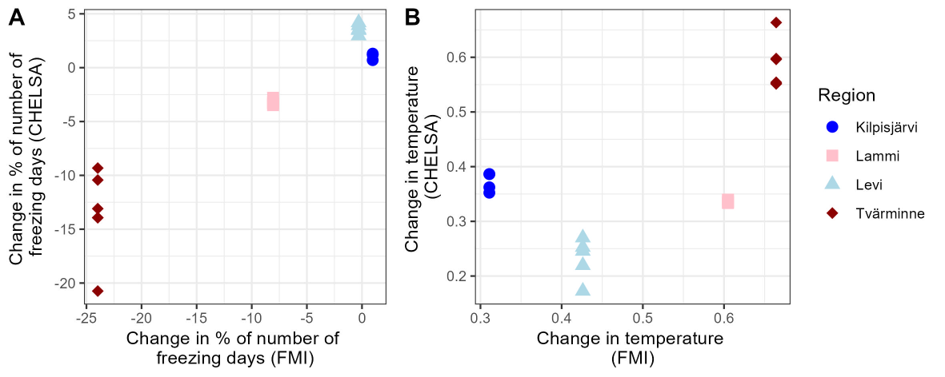

Figure S2: Comparison between climate change data from CHELSA database and meteorological stations of Finnish Meteorological Institute (FMI). A: Comparison of the change in % of the number of days in the year when mean daily temperature is <0°C, between 2005-2014 and 2015-2024 time periods. B: Comparison of mean daily temperature change, between 2005-2014 and 2015-2024 time periods. Data was extracted for each site with the exact GPS coordinates for CHELSA data, and for the nearest meteorological stations for FMI (same as described in Table S1, one station for each region). Dark blue circles, light blue triangles, pink squares, and dark red diamonds correspond to sites from, respectively, Kilpisjärvi, Levi, Lammi, and Tvärminne regions.

Table S2: Abundance per site and occupancy of each species in the initial sampling (2015) and the resampling (2025-2026). Abundance is given as mean ± standard deviation, occupancy is given as the number of sites in which the species is present, *i.e.,* the sum of occurrences. In each site 0.5625 m² were sampled (0.375 m² for litter sorting + mustard extraction, 0.1875 m² for litter and soil hand sorting). Results are shown for all individuals and for adults only. P-values are permutational p-values, those of the one-tailed paired t-test with 999 permutations.

|  | Abundance | |  | Occupancy | |
| --- | --- | --- | --- | --- | --- |
|  | 2015 | 2025-2026 | P-value | 2015 | 2025-2026 |
| *Aporrectodea caliginosa* |  |  |  |  |  |
| all individuals | 2.00±3.92 | 3.14±6.26 | 0.150 | 7 | 6 |
| adults only | 2.00±3.92 | 2.19±4.80 | 0.480 | 7 | 6 |
| *Aporrectodea rosea* |  |  |  |  |  |
| all individuals | 1.19±4.38 | 3.43±1.00 | 0.127 | 3 | 3 |
| adults only | 1.19±4.38 | 2.05±5.74 | 0.253 | 3 | 3 |
| *Bimastos rubidus* |  |  |  |  |  |
| all individuals | *0.14±0.36* | *0.90±2.84* | *0.086* | 3 | 5 |
| adults only | 0.10±0.30 | 0.24±0.54 | 0.261 | 2 | 4 |
| *Dendrobaena octaedra* |  |  |  |  |  |
| all individuals | **11.38±15.77** | **6.05±5.77** | **0.014** | 16 | 18 |
| adults only | **1.76±2.45** | **0.71±1.01** | **0.032** | 12 | 10 |
| *Lumbricus castaneus* |  |  |  |  |  |
| all individuals | *not found* | 0.10±0.44 | 0.499 | 0 | 1 |
| adults only | *not found* | 0.10±0.44 | 0.505 | 0 | 1 |
| *Lumbricus rubellus* |  |  |  |  |  |
| all individuals | 0.24±0.89 | 0.19±0.51 | 0.491 | 2 | 3 |
| adults only | 0.24±0.89 | 0.10±0.30 | 0.513 | 2 | 2 |
| *Lumbricus terrestris* |  |  |  |  |  |
| all individuals | 1.10±2.76 | 0.90±1.64 | 0.394 | 5 | 6 |
| adults only | 1.10±2.76 | 0.81±1.44 | 0.324 | 5 | 6 |
| *Octolasion lacteum* |  |  |  |  |  |
| all individuals | 0.10±0.30 | 0.19±0.87 | 0.524 | 2 | 1 |
| adults only | 0.10±0.30 | 0.14±0.65 | 0.533 | 2 | 1 |

Table S3: Adult earthworm abundance and species richness per site in the initial sampling (2015) and the resampling (2025-2026). Adults were considered as individuals with a clitellum. In each site 0.5625 m² were sampled (0.375 m² for litter sorting + mustard extraction, 0.1875 m² for litter and soil hand sorting). Results are showed as mean ± standard deviation. The table shows mean site abundance and richness for all sites and among each region. ^a^ P-value of Wilcoxon signed rank test = 0.05791. All other p-value of Wilcoxon signed rank test > 0.1.

|  | Abundance | | Richness | |
| --- | --- | --- | --- | --- |
|  | 2015 | 2025-2026 | 2015 | 2025-2026 |
| All sites (n=21) | 6.48±2.14 | 6.33±2.16 | 1.57±0.40 | 1.57±0.38 |
| Kilpisjärvi (n=4) | 0.25±0.25 | 0.75±0.48 | 0.25±0.25 | 0.50±0.29 |
| Levi (n=5) | 0.40±0.40 | 0.40±0.40 | 0.40±0.40 | 0.20±0.20 |
| Lammi (n=6) | 10.83±5.39 | 14.00±4.80 | 2.50±0.92 | 3.20±0.75 |
| Tvärminne (n=6) | 11.33±3.86^a^ | 7.33±4.45^a^ | 2.50±0.67 | 1.83±0.60 |
